# Controlling the taxonomic composition of biological information storage in 16S ribosomal RNA

**DOI:** 10.1101/2025.04.29.651329

**Authors:** Kiara Reyes Gamas, Travis R. Seamons, Matthew J. Dysart, Lin Fang, James Chappell, Lauren B. Stadler, Jonathan J. Silberg

## Abstract

Microbes can be programmed to record participation in gene transfer by coding biological-recording devices into mobile DNA. Upon DNA uptake, these devices transcribe a catalytic RNA (cat-RNA) that binds to conserved sequences within ribosomal RNA (rRNA) and perform a trans-splicing reaction that adds a barcode to the rRNA. Existing cat-RNA designs were generated to be broad-host range, providing no control over the organisms that were barcoded. To achieve control over the organisms barcoded by cat-RNA, we created a program called Ribodesigner that uses input sets of rRNA sequences to create designs with varying specificities. We show how this algorithm can be used to identify designs that enable kingdom-wide barcoding, or selective barcoding of specific taxonomic groups within a kingdom. We use Ribodesigner to create cat-RNA designs that target Pseudomonadales while avoiding Enterobacterales, and we compare the performance of one design to a cat-RNA that was previously found to be broad host range. When conjugated into a mixture of *Escherichia coli* and *Pseudomonas putida*, the new design presents increased selectivity compared to a broad host range cat-RNA. Ribodesigner is expected to aid in developing cat-RNA that store information within user-defined sets of microbes in environmental communities for gene transfer studies.

**GRAPHICAL ABSTRACT:** 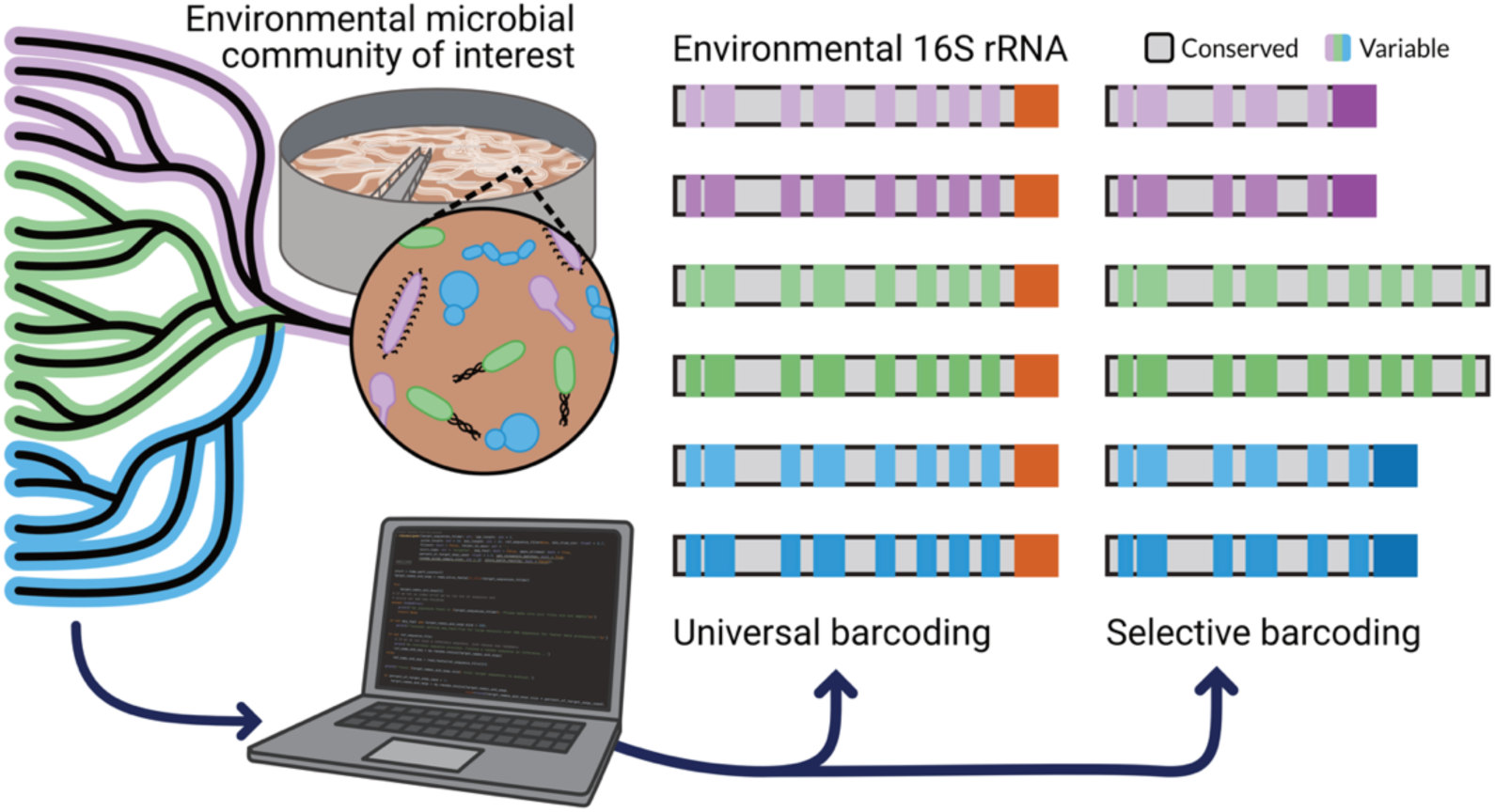

## INTRODUCTION

Biosensors have the potential to transform our understanding of environmental processes,^1,2^ since they can couple the detection of specific environmental conditions to the production of easy-to-detect output signals.^3–5^ Broadly, there are two classes of microbial sensors. First, real-time biosensors provide continuous information on environmental chemicals, generating visual or chemical outputs continuously when they encounter the inputs.^6–9^ Second, memory-based biosensors record and store information about sensing using mechanisms that can be read out at a later point in time.^10–13^ These latter biosensors have advantages with monitoring transient processes within environmental communities, such as gut,^14,15^ soil,^16,17^ and wastewater^18^ communities. Further, memory biosensors can record information about complex biological processes like gene transfer,^19^ which underlies the spread of antibiotic resistance^20,21^ and is critical to understand when engineering microbes for environmental applications.^22^

Memory biosensors can record information in nucleic acids using either permanent sequence modifications,^23–25^ or using mechanisms that enable the system to be reset.^26,27^ A variety of genetic memory systems have been developed to support studies of gene transfer. Notably, CRISPR recording tapes being have been created that sense gene transfer between a gut microbial community and an engineered *Escherichia coli*.^14^ More recently, mobile genetic elements built using the *mariner* transposon have been used to record gene transfer in a microbial community.^19^ With this latter approach for studying gene transfer, mobile DNA host range is obtained through metagenomic sequencing, like high-throughput chromosome conformation capture.^28^ While these approaches can store information across diverse species within microbial communities, all genetic material in an environmental sample must be sequenced to read out stored information.

Recently, a catalytic RNA (cat-RNA) was described that can be coded into mobile DNA and used to record information about gene transfer host range.^29^ As shown in Figure 1a, this cat-RNA, which is built using the *Tetrahymena* group I intron splicing ribozyme,^30,31^ splices an RNA barcode onto 16S ribosomal RNA (rRNA) adjacent to variable regions used for taxonomic identification. Targeted sequencing of barcoded 16S rRNA can then be used to establish which microbes in a community participated in gene transfer. Proof-of-concept measurements with this RNA addressable modification (RAM) device revealed that cat-RNA can record community-level information about participation in gene transfer within diverse gram-negative bacteria. While the prototype cat-RNA achieved broad host range activity in a wastewater community, it was designed using an alignment of five 16S rRNA. As such, there is a need for a more rigorous design approach for cat-RNA.

**Figure 1.**
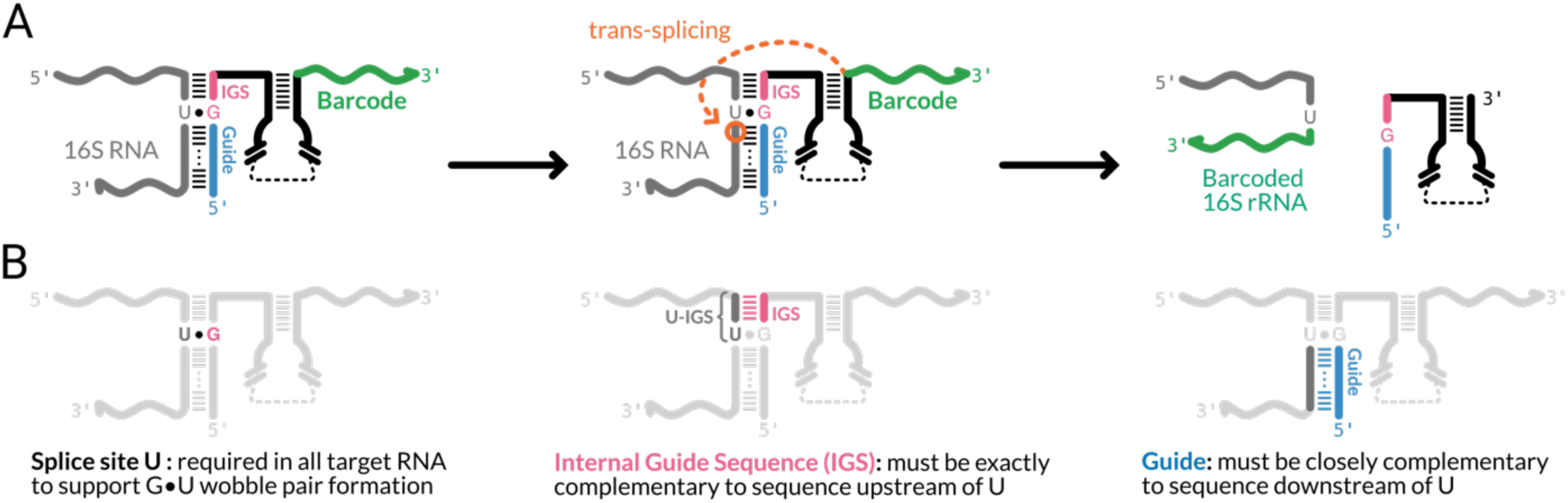
Cat-RNA design considerations for 16S rRNA recording. (**A**) Cat-RNA have a G that is flanked by a guide (blue) and IGS (pink), as well as a barcode (green) that is ultimately amended to 16S rRNA. When the guide and IGS anneal to 16S rRNA, the G forms a G·U wobble pair with the U targeted for splicing, and the barcode gets spliced downstream of this site. (**B**) Ribodesigner makes three assumptions: (i) a uracil substrate is critical for the catalytic reaction, (ii) the splicing efficiency depends upon the IGS being complementary across all five base pairs to the sequence upstream of the targeted U, and (iii) the guide must anneal to the 16S rRNA downstream of the U targeted for splicing.

Herein, we describe a computational program called Ribodesigner that identifies cat-RNA designs that either maximize information storage across a set of rRNA sequences or selectively target specific groups within the set. Ribodesigner leverages design principles that have been successful in primer design software.^32–34^ We use Ribodesigner to identify universal cat-RNA that are expected to maximize information storage across Archaea, Bacteria, or Eukaryotes. We also show how Ribodesigner can be used to design cat-RNA for selectively storing information in one order while avoiding a second order. Finally, we compare a pair of designs that are predicted to vary in their barcoding of Pseudomonadales and Enterobacterales and observe the predicted variation in barcoding selectivity.

## RESULTS

### Designing universal cat-RNA

We hypothesized that three design parameters would be important for programming cat-RNA to barcode rRNA across diverse taxonomic groups (Figure 1b). First, we hypothesized that the target 16S rRNA must contain an uracil at the targeted splicing site to support the formation of a G•U wobble and catalysis.^35,36^ Second, to maximize catalytic efficiency, we posited that the pentanucleotide sequence upstream of the U splice site must be complementary to the internal guide sequence (IGS) in the synthetic ribozyme across all five base pairs.^37^ Prior studies of trans-splicing ribozymes have revealed that splicing activity depends upon the formation of a base-paired helix called P1 between the IGS and sequences adjacent to the splice site.^38,39^ Third, we assumed that the ribozyme must have a guide sequence that directs annealing to the 16S rRNA sequence downstream of the U splice site to support cat-RNA and 16S rRNA association at the desired location. The guide was included as a requirement since a recent study showed that split ribozymes can be spliced together when they are designed to anneal to a template RNA.^40^ All guide designs were 50 base pairs long, because this length presented the highest efficiency in the original RAM design.^29^

To design cat-RNAs for recording information in communities, Ribodesigner identifies barcoding sites in user-defined sets of rRNA through four steps. *<u>First</u>*, the user provides a set of rRNA sequences for creating designs, defined as the ‘design set’, and a reference rRNA sequence for indexing all sequences during the analysis. The user selects the guide length and a minimum level of complementarity desired in the cat-RNA designs. *<u>Second</u>*, the algorithm aligns each unique rRNA sequence in the design set to the reference sequence and scores U frequencies at every position. *<u>Third</u>*, the algorithm assesses the frequencies of different pentanucleotide sequences upstream of each U in the rRNA set, which are designated U-IGS sequences. These sequences will ultimately serve as the IGS-binding sites for the cat-RNA. *<u>Fourth</u>*, the algorithm creates guide sequences for every unique U-IGS and scores them based on complementarity to a second set of rRNA sequences, defined as the ‘evaluation set’.

To understand the conservation of uracils in rRNA and adjacent IGS-binding pentanucleotide sequences across bacteria, we applied Ribodesigner to a set of 2410 bacterial 16S rRNA (Figure 2a). This design set, which came from 834 bacterial orders, was chosen to capture the breadth of bacterial diversity in the SILVA rRNA database^41^ and minimize degeneracy of closely-related species. Across the sequence space of this bacterial 16S rRNA design set, a wide range of U frequencies were observed. Highly conserved uracils, defined as sites found in >75% of test sequences, were dispersed across the 16S rRNA sequence (Figure 2b). Within the 16S rRNA variable regions, uracils were more frequently flanked by IGS-binding motifs that varied in sequence (Figure 2c), while uracils between the variable regions had IGS-binding motifs with the highest conservation (Figure 2d). Across the 2410 bacterial rRNA, 44 of the positions had U-IGS with >75% identity. Similar trends were observed when Ribodesigner was applied to sets of 337 Archaeal 16S rRNA (Figure S1) and 2618 Eukaryotic 18S rRNA (Figure S2). The optimal designs observed when targeting each kingdom differed. Thus, Ribodesigner can identify highly conserved U-IGS when provided different sets of rRNA sequences.

**Figure 2.**
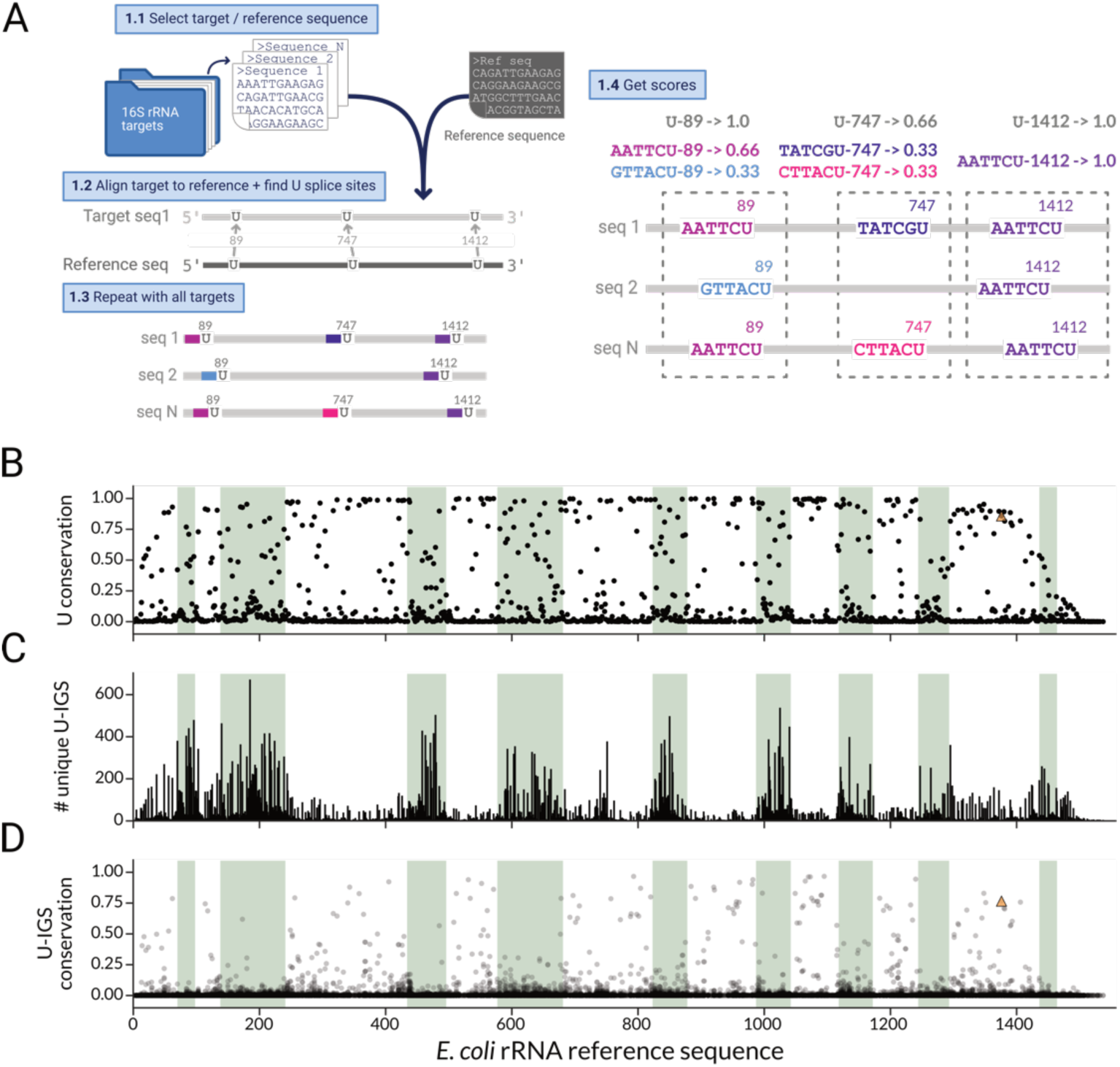
Uracil conservation across bacterial 16S rRNA. (**A**) The workflow that Ribodesigner uses to calculate: (**B**) the conservation of U at each 16S rRNA position in the design set, (**C**) the diversity of IGS-binding pentanucleotide sequences adjacent to each U (U-IGS), and (**D**) the conservation of each U-IGS across the design set of 16S rRNA. The bacterial 16S rRNA design set included 2410 rRNA sequences. The hypervariable regions (green) are indexed to the reference sequence, *E. coli* MG1655. The previously characterized broad-host range cat-RNA, which targets U1376 in *E. coli* rRNA, is shown as an orange triangle.^29^

To design cat-RNA guides that maximize annealing, the 16S rRNA sequences downstream of each U-IGS are analyzed. This step is performed by generating guide sequences adjacent to each U-IGS using small batches (n = 10) of rRNA sequences from the design set and then generating a consensus sequence for each of those batches (Figure 3a). For each unique U-IGS, the consensus sequence with the lowest ambiguity (Figure S3) is identified by scoring it through alignment with 16S rRNA in the bacterial evaluation set (Figure 3b), which is distinct from the set used for design. To further improve the annealing scores of the guides, the guide sequences with the lowest ambiguities from each batch are pooled and used to find consensus sequences using the same process (Figure 3c). When the fifty designs having the highest U-IGS scores were mapped onto the 16S rRNA sequence (Figure 3d), a subset of designs were found to have high guide scores (Figure 3e), which were dispersed across the different conserved motifs. The cat-RNA previously designed for broad host range barcoding using five rRNA sequences was among the top designs,^29^ although it did not present the highest IGS and guide scores. Similar high-quality cat-RNA designs could be identified when performing this analysis with sets of 337 Archaeal rRNA (Figure S4) and 2618 Eukaryotic rRNA (Figure S5). The optimal designs targeting each kingdom were different. These analyses show how Ribodesigner can be used to identify guide designs that are predicted to support binding to diverse rRNA adjacent to highly conserved U-IGS within user-defined sets of rRNA. In addition, they suggest that different cat-RNA designs should be used to achieve broad-host range barcoding of rRNA in organisms from different kingdoms.

**Figure 3.**
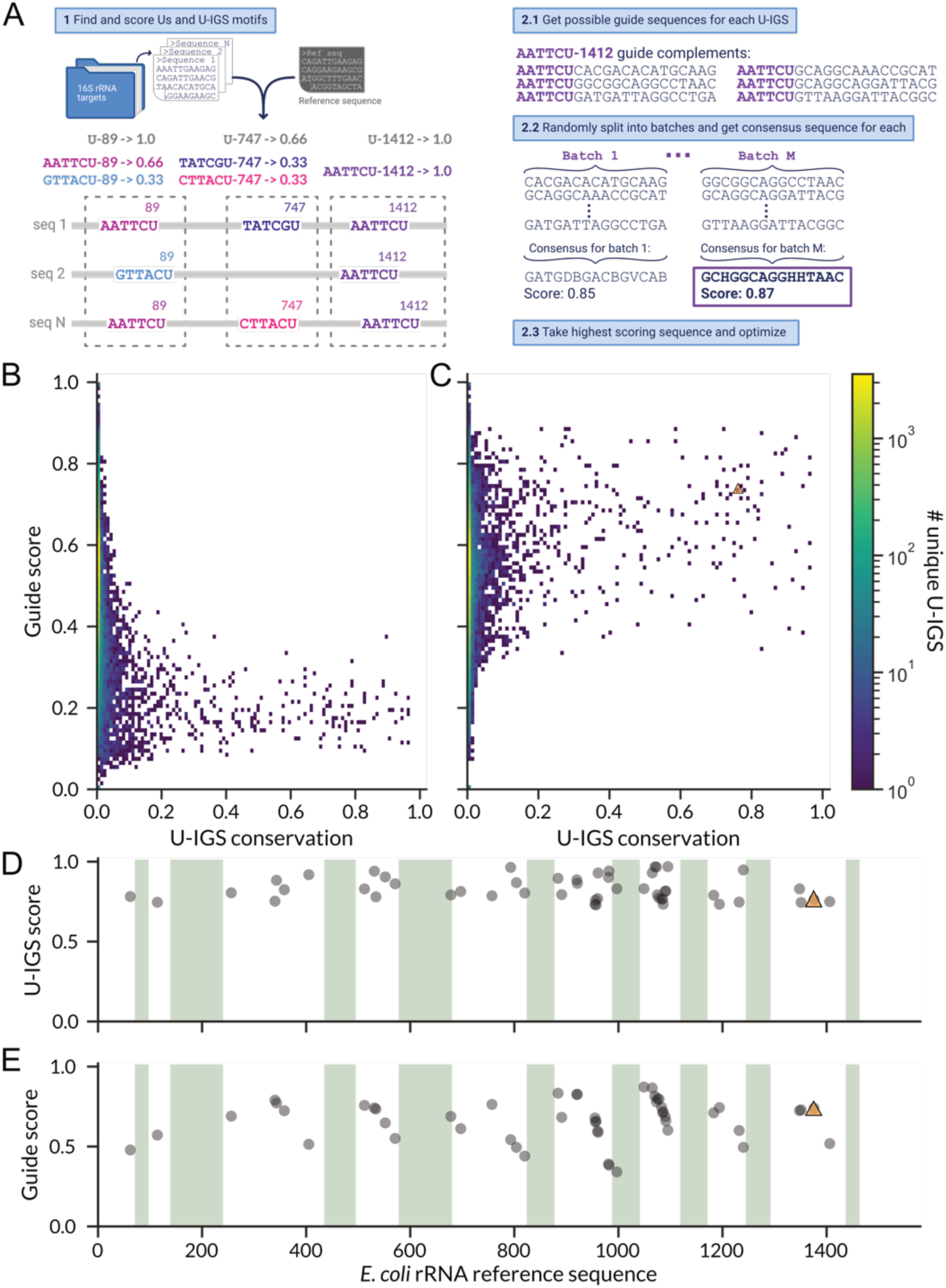
Designing cat-RNA guide sequences for bacteria. (**A**) For every unique U-IGS, guides are created by generating consensus sequences using small batches of 16S rRNA sequences upstream of each targeted U (n = 10). Using a design set of 2410 16S rRNA, guides were designed against all 65155 unique U-IGS motifs. (**B**) The U-IGS conservation and guide scores for all 65155 cat-RNA generated using the evaluation set of 16S rRNA. (**C**) Guide score improvements achieved by using the winning guides from each batch to create optimized guides. (**D**) For the 50 designs having the highest U-IGS scores, the location of U targeted for barcoding is mapped onto *E. coli* 16S rRNA. (**E**) For those same designs, the guide scores are also mapped onto *E. coli* 16S rRNA. The cat-RNA previously applied in a wastewater is shown as an orange triangle.^29^

### Creating selective cat-RNA

The previously characterized cat-RNA exhibits broad host range in communities.^29^ However, there may be cases where it is desirable to program cat-RNA to more selectively record information about gene transfer in a subset of community members that take up mobile DNA. To design cat-RNA with narrower taxonomic selectivity, we created an option in Ribodesigner to generate a selectivity score (Figure 4a). For this analysis, three sets of rRNA must be designated, including: (i) a design set of rRNA targeted for barcoding, (ii) an off-target rRNA evaluation set that are to be avoided, and (iii) an on-target evaluation set that differs from the design set. First, using the design set, cat-RNA are generated and scored. Second, the designs from this analysis are scored independently using the on- and off-target evaluation sets. Finally, a selectivity score is calculated for each design by subtracting the off-target score from the on-target score. With this approach, scores having a value of one indicate high selectivity for the target rRNA set, while lower scores indicate poorer selectivity.

**Figure 4.**
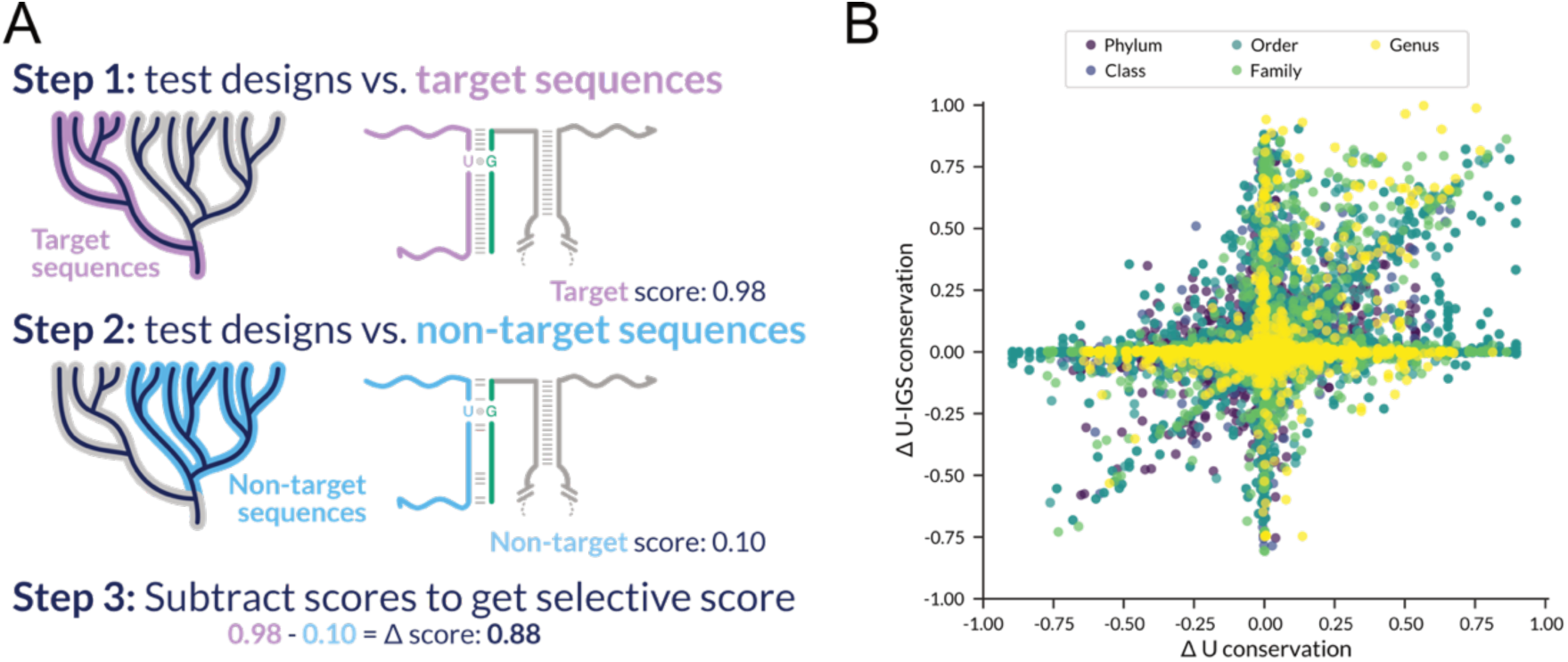
Evaluating the selectivity of cat-RNA designs at different taxonomic levels. (**A**) To estimate cat-RNA selectivity, Ribodesigner separately scores the cat-RNA components against a subset of the community (target sequences) and all other members of the community (non-target sequences). The program then subtracts the scores obtained for each, such that the difference indicates the selectivity for the target organisms. (**B**) For an *in silico* community, a specificity score for each U (ΔU conservation) and each U-IGS (ΔU-IGS conservation) was calculated for each taxonomic group at the phyla, class, order, family, and genera level.

To evaluate the extent to which microbes from different taxonomic groups can be differentiated using the selectivity score, we designed taxonomically-selective cat-RNA using microbes commonly found in wastewater microbiomes. For these calculations, we constructed an *in silico* community made up of microbes from three phyla, including gram-negative (*Proteobacteria*) and gram-positive (*Actinobacteriota* and *Firmicutes*) bacteria that are often abundant in wastewater.^29^ This *in silico* community was assembled by creating a set of 16S rRNA sequences from the SILVA database.^41^ To determine the extent to which cat-RNA can be selectively designed for each taxonomic group in the community, we calculated: (i) the conservation of every U and U-IGS for different target groups in this community, (ii) the conservation of U and U-IGS at the same sites in all other organisms in the community, which represent non-target groups, and (iii) the differences between these scores for uracils (ΔU score) and U-IGS (ΔU-IGS score). A wide range of selectivity scores were observed with this analysis (Figure 4b). Among these, scores indicative of high selectivity (>0.75) were observed when evaluating designs targeting a subset of organisms for barcoding at the genus, family, and order levels. These findings suggest that it may be possible to achieve barcoding selectivity among different organisms at the genus, family, and order levels by targeting U for splicing that present high ΔU and ΔU-IGS scores.

The selectivity calculations with Ribodesigner suggested that cat-RNA designs can be created that barcode 16S rRNA in some organisms while avoiding barcoding in other organisms. To test this idea, we used Ribodesigner to identify cat-RNA designs that are predicted to selectively barcode Pseudomonadales while minimizing barcoding of Enterobacterales. These calculations were performed using 3000 16S rRNA sequences from each order, which were compiled using the SILVA database.^41^ The Pseudomonadales design with the highest calculated selectivity (Pcat-RNA), which is predicted to perform a trans-splicing reaction after variable region six within 16S rRNA, was compared with a broad-host range, universal cat-RNA design previously characterized in a wastewater community. When evaluating both designs with the same set of Enterobacterales used for design, the U and U-IGS scores were larger for the universal cat-RNA compared to Pcat-RNA (Figures. 5a-b). In contrast, when evaluating these designs with against the set of Pseudomonadales used for design, similar high U and U-IGS scores were obtained for both the universal cat-RNA and Pcat-RNA. Both the universal cat-RNA and Pcat-RNA presented high guide scores with both target and non-target organisms (Figure 5c). These findings identify a Pcat-RNA design that is predicted to present distinct barcoding selectivity compared to the broad-host range cat-RNA, selectively barcoding 16S rRNA in Pseudomonadales but not in Enterobacterales.

**Figure 5.**
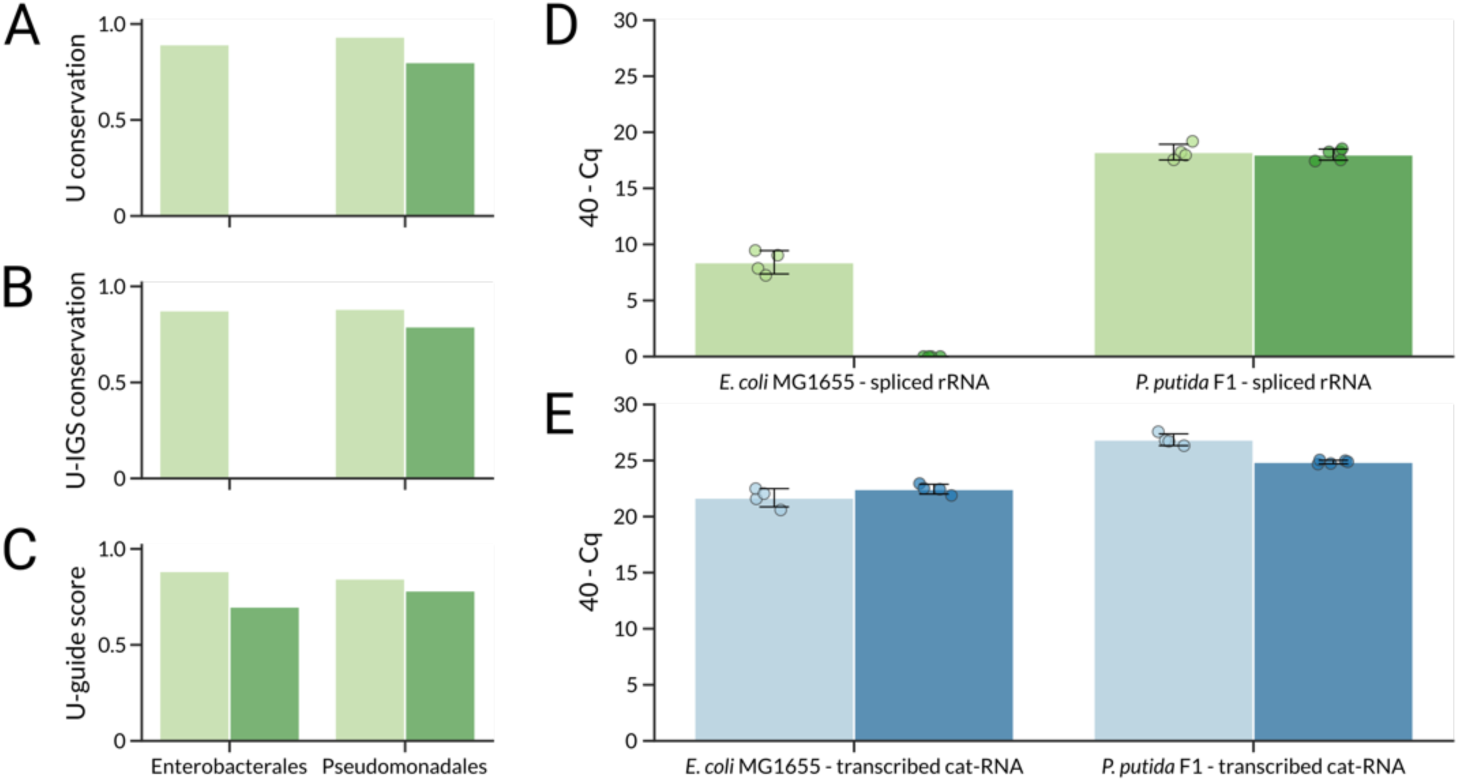
Comparing the selectivity of different cat-RNA designs. *In silico* comparison of the (**A**) U, (**B**) U-IGS, and (**C**) guide scores for Pcat-RNA (dark green) and universal cat-RNA (light green). Values were calculated using evaluation sets of 3000 Enterobacterales and 3000 Pseudomonadales. (**D**) The splicing efficiency of Pcat-RNA was compared with the universal cat-RNA using qPCR in model organisms from the targeted order (*P. putida* F1) and non-targeted order (*E. coli* MG1655). With *E. coli* MG1655, the qPCR signal was below the limit of detection. (**E**) Total transcribed cat-RNA was quantified using qPCR in cells transcribing the universal cat-RNA (light blue) and Pcat-RNA (dark blue). For each experiment, error bars represent ±10 from four biological replicates were performed.

We next created a plasmid that transcribes the Pcat-RNA by altering the IGS and guide sequence in the previously described broad host range cat-RNA.^29^ To evaluate if Pcat-RNA selectively targets *Pseudomonadales*, the plasmid was transformed into *E. coli* MG1655 and *Pseudomonas putida* F1, respectively. Strains were also transformed with a previously described broad host range cat-RNA (“universal cat-RNA”) as a control.^29^ Cells were grown overnight, total RNA was isolated, and total cat-RNA and barcoded 16S rRNA were quantified using reverse-transcription quantitative PCR (RT-qPCR). With this analysis, the universal cat-RNA yielded a signal for barcoded-rRNA in both *E. coli* MG1655 and *P. putida* F1 (Figure 5d). In contrast, Pcat-RNA only presented a barcoded-rRNA signal in *P. putida* F1. No signal was observed in *E. coli* MG1655. As a control, we evaluated the level of transcribed ribozymes in *E. coli* MG1655 and *P. putida* F1. Figure 5e shows that cat-RNA and Pcat-RNA were transcribed at similar levels in each organism. Further, similar levels of total 16S rRNA were observed in each of the samples (Figure S6). Together these findings demonstrate evidence of selectivity, as Pcat-RNA yielded a barcoding signal in Pseudomonadales but not in Enterobacterales, despite being transcribed to a similar extent as the universal cat-RNA.

To investigate if next-generation sequencing (NGS) can detect selective barcoding in a gene transfer experiment, we transformed *E. coli* MFDpir^42^ with conjugative plasmids that transcribe Pcat-RNA and the universal cat-RNA and added each strain to a community containing *E. coli* MG1655 and *P. putida* KT2440 at similar titers (Figures 6a-b). Following conjugation for 24 hours, total RNA was isolated from each sample and converted into complementary DNA (cDNA), and amplicon sequencing was performed (Figures S7-S8). With the universal cat-RNA, a comparison of the ratios of barcoded rRNA to native 16S rRNA read counts (Figure 6c) revealed that the signal in *E. coli* MG1655 was 9-fold higher than in *P. putida* KT2440. In contrast, Pcat-RNA presented a signal in *P. putida* KT2440 that was 26-fold higher than the ratio of barcoded rRNA to native 16S rRNA in *E. coli* MG1655. Only the signals with the Pcat-RNA were significantly different across the strains. These findings demonstrate that Ribodesigner conjugation of selective cat-RNA can store information in a targeted subset of community members.

**Figure 6.**
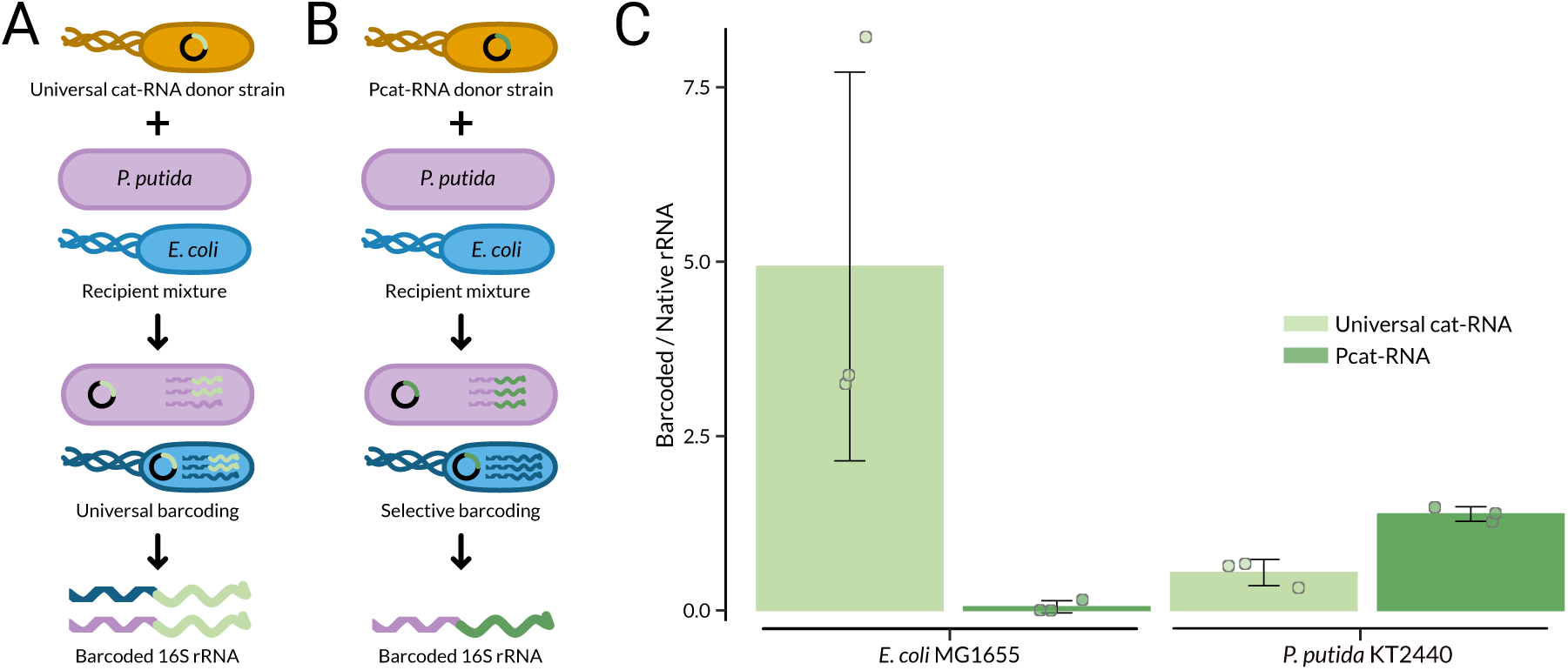
Cat-RNA selectivity achieved in a synthetic community. *E. coli* MFDpir was used to conjugate either (**A**) a broad host range cat-RNA (universal) and (**B**) a cat-RNA designed to be Pseudomonadales selective (Pcat-RNA) into a two-member community containing *E. coli* MG1655 and *P. putida* KT2440. Following conjugation, both native and barcoded 16S rRNA were sequenced. (**C**) Barcoded-rRNA counts normalized to 16S rRNA for the universal (light green) and Pcat-RNA (dark green). Each bar represents the ratio between rarefied counts (n = 100000), while error bars are ±10 from three biological replicates. The Pcat-RNA signal is significantly lower in *E. coli* MG1655 compared to *P. putida* KT2440 (Welch’s two tailed, unpaired t-test, p = 0.006), while the universal cat-RNA was not significantly different across the strains (p = 0.1).

### Implications

Ribodesigner represents a new computational tool for designing cat-RNA with user-defined barcoding host ranges. By calculating a selectivity score, we show how Ribodesigner can identify cat-RNA designs that are predicted to selectively barcode microbes in the order Pseudomonadales while avoiding a signal in Enterobacterales. We validate one selective cat-RNA design by showing that it can be used for conjugation in a synthetic community to increase the barcoded-rRNA signal in a targeted microbe compared with a previously described broad-host range cat-RNA. In future studies, it will be interesting to investigate how this selective cat-RNA performs in communities containing a greater diversity of Enterobacterales and Pseudomonadales, such as wastewater samples.^43,44^ Selective cat-RNA designs are expected to improve RAM’s ability to report on gene transfer within low-abundance microbes in communities, which will be important for studies of engineered bacteria for environmental release.^45^ By enabling the selective barcoding of rRNA in a subset of organisms that participate in gene transfer, a greater fraction of the barcoded-rRNA sequencing reads is expected to represent the targeted organisms, similar to increases in sensitivity that are achieved with hybridization capture enrichment approaches.^46^ Ribodesigner will also be useful for designing broad-host range cat-RNA for monitoring gene transfer within the different kingdoms of life. The archaeal designs should be useful for studying the host range of liposome-mediated transformation methods commonly used with isolates by applying them in communities rich in archaea,^47,48^ while the eukaryote designs can be used to study the host range of Agrobacterium-mediated transformation methods and horizontal gene transfer in fungi.^49,50^

## METHODS

### Calculations

Ribodesigner, which can be found on Github (kiararey/Ribodesigner), takes in user-defined sets of small subunit (SSU) rRNA sequences and finds cat-RNA designs for all possible U splicing sites in each sequence. When creating broad-host range designs, the algorithm performs five different calculations. First, the algorithm aligns each target rRNA sequence in the ‘design set’ to a reference sequence, which is defined by the user, and indexes each potential splice site. Second, all potential splice sites in the design set are scored by calculating the fraction of sequences in the set with a conserved U at the same location. Third, the algorithm calculates the conservation of the five base pairs adjacent to each U, which will ultimately bind to the IGS in the cat-RNA. In some cases, a given site in the rRNA has multiple U-IGS, which differ in their frequencies. Third, using the subset of rRNA sequences having conserved U-IGS sequences at a given site, Ribodesigner extracts every possible guide sequence that would match the 50 bases downstream of the conserved U. It uses these sequences to calculate a consensus sequence that maximizes the base pairing conservation across all target sequences. Fourth, the program scores each of these guide sequences based on their ambiguity. Fifth, the algorithm scores each cat-RNA design against an evaluation set of rRNA sequences, defined by the user which is distinct from the design set. To score quality, three parameters are calculated for every location in the indexed rRNAs, including: (i) the fraction of rRNA sequences containing a U, (ii) the fraction of rRNA sequences containing each observed U-IGS sequence, and (iii) the average correct base pairing of the guide to the test sequences. To create taxa-specific designs, Ribodesigner uses the aforementioned protocol with a design set of rRNA sequences. Then, it scores the designs generated against an ‘on-target evaluation set’ of rRNA where barcoding is desired and an ‘off-target evaluation set’ where barcoding is to be avoided. Finally, for each cat-RNA design, it subtracts the score from the off-target evaluation from the on-target evaluation to estimate the selectivity.

<u>Step 1: Alignment of target sequences to a reference sequence.</u> The input to the ribodesigner.py program is a FASTA file containing a design set of target SSU sequences. These sequences are aligned to a reference sequence, defined by the user, using a global alignment with Bio.Align.PairwiseAligner in Biopython. The user also defines the reference sequences and SSU rRNA variable regions. For this study, the SSU rRNA sequences for bacteria (*Escherichia coli* MG1655), eukaryotes (*Saccharomyces cerevisiae*), and archaea (*Methanobrevibacter smithii*) differed. Variable regions were labeled for *E. coli* and *S. cerevisiae* based on literature,^51,52^ while the *M. smithii* variable regions were determined by finding conserved regions with a pairwise alignment with the *E. coli* 16S rRNA sequence using the function adjust\_var\_regs in Ribodesigner.

<u>Step 2: Identification and evaluation of U-IGS sequences.</u> The program finds all potential U splice sites, defined as all uracils in each rRNA sequence that are at least six nucleotides downstream from the 5’ end and at least 50 nucleotides away from the 3’ end. The latter constraint is longer because there must be sufficient sequence for cat-RNA guide annealing. Each U site is given an index corresponding to the position of that base pair in the reference sequence. The program then finds the corresponding IGS and guide sequences that a cat-RNA would require to target each potential U splice site, defined as the reverse complement of the five bases upstream of the U splice site and the reverse complement of the 50 bases downstream of the U splice site, respectively. This yields a variety of cat-RNA designs, each consisting of a guide sequence at the 5’ end fused to a G and an IGS. Ribodesigner aligns conserved SSU rRNA sequences, so the computational time generating adequate putative sequences can be drastically reduced by first aligning each target sequence to a shared reference sequence and finding the equivalent reference position (reference index) for each potential U splice site. Ribodesigner thus does a pairwise alignment between a given reference sequence and each target sequence using a global alignment with Bio.Align.PairwiseAligner. Ribodesigner scores each design by calculating the U-IGS true coverage, defined as the fraction of all sequences in the design set with that exact U-IGS sequence at the same reference index position.

<u>Step 3: Guide sequence generation.</u> Once all possible U-IGS have been identified and scored in the design set of rRNA, the program extracts and pools together the corresponding complementary guide sequences, which will ultimately be used to build cat-RNA. These guides are randomly grouped into batches of 10. The guide sequences in each batch are then aligned with MUSCLE5 multiple sequence alignment.^53^ A consensus sequence is generated for each batch. To generate the consensus sequence, we modified the consensus sequence generation function in RSAT to include ambiguity nucleotide codes.^54^ Briefly, a list of sequences are fed into the “words2countmatrix” function, as well as a priori containing the base frequency of each base. In this study, the priori was kept as 0.25 for either A, T, G, C. This function counts the base of each sequence and adds up the frequency of each base at each position, returning a matrix with base frequency counts at each position for the input set of sequences. Then, the function “consensus” counts the number of bases at each position and returns the most likely base. If there are equal probabilities of more than one base at one position, the corresponding ambiguity code is returned. For example, if both A and G have equal likelihood in one position, the code “R” is returned.

<u>Step 4: Guide sequence optimization.</u> Each batch of guides results in one consensus guide sequence. Each consensus guide sequence is scored using the weighted scoring method described in the appendix, and the highest-scoring sequence is selected as the template design for the guide. If the template contains ambiguous bases, an additional optimization protocol is performed to reduce ambiguity. Ambiguity is reduced by performing pairwise alignments between the template and each of the other batch consensus guide sequences, in order of highest to lowest score. For each pairwise alignment, at each location in the template with an ambiguous base, the corresponding base at the same location in the other sequence is compared. If the corresponding base can reduce the ambiguity (e.g., contains a shared base), then the base in the template design is replaced by the shared base. For example, if a base in the template design is ‘H’, corresponding to bases ‘A’, ‘C’, or ‘T’, and the corresponding base in the other sequence is a ‘T’, the base in the template design is replaced by the shared base ‘T’. This process repeats until either the guide score increases to 1, includes no ambiguous bases, or until the consensus sequence of each batch has been compared. For a guide sequence *x* with *n* total bases and length of *m*, the weighted score (*S*_*weighted*_) is defined as:

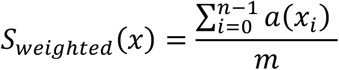

where *a(x_i_)* is the degenerate score for the base *x* at location *I* such that:

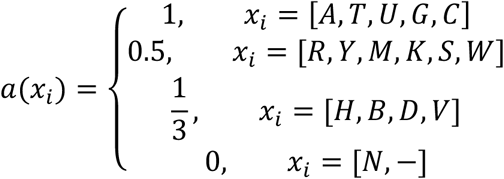

<u>Step 5. Design evaluation using test sequences.</u> The quality of broad host range cat-RNA designs is scored using an evaluation set of rRNA. With this evaluation, three aspects of the cat-RNA are scored, including: (i) the conservation of the U targeted in the rRNA sequences in the evaluation set, (ii) the conservation of the U-IGS sequence at the same location in the evaluation set, and (iii) the average guide conservation (guide score). The U conservation of each design is defined as the fraction of test sequences containing a U at the same index targeted by the design. The U-IGS conservation is defined as the true coverage of the design’s U-IGS sequence in the test sequences with a U splice site at the same index targeted by the design. Finally, the guide score is defined as the mean of the weighted guide score of the consensus sequence arising from the alignment between the design’s guide sequence and each test sequence. The quality of selective cat-RNA designs is scored using a pair of evaluation sets of rRNA, including on- and off-target sets. Each of these scores is calculated as described for the broad host range designs. Then the specificity score is generated by subtracting the off-target score from the on-target score.

### RNA sequences used for calculations

Table S1 provides a summary of the datasets used for Ribodesigner calculations, which were generated from the SILVA SSU Ref NR99 database. These data sets can be found on Github (github.com/kiararey/Ribodesigner/tree/main/Common_sequences). These data sets fall into three categories, including: (i) three design sets of target rRNA sequences from each kingdom used for creating broad-host range designs, (ii) a set of microbes whose diversity mirrors that in a wastewater community for selectivity calculations, and (iii) sets of Enterobacterales and Pseudomonadales used for designing and testing the Pcat-RNA. The protocols for selecting these data sets are described below.

<u>Broad-host range designs.</u> To test whether the program could make universal designs, we needed a dataset that would be representative of a large variety of organisms while avoiding bias arising from members overrepresented in the database. To do so, we downloaded the SILVA Ref Seq SSU NR 99 dataset and chose a subset.^41^ For universal design, we randomly selected 10 organisms per genus from each kingdom that did not belong to the same species. One of these datasets was used to make cat-RNA (design set) while the other was used to test designs (evaluation set). If there were fewer than 10 unique genera in an order, the design and evaluation sequences contained overlapping sequences. This resulted in 2 reduced SSU datasets for each kingdom.

<u>Wastewater designs.</u> To evaluate how well selectivity can be achieved, we made datasets corresponding to different taxonomic classification levels of the top three orders observed in wastewater from a previous RAM study,^29^ which included Aeromonadales, Enterobacterales, and Pseudomonadales, as well as gram-positive organisms. In the SILVA database, the orders Aeromonadales and Enterobacterales are both classified as Enterobacterales. These datasets included 10 sequences per genus. We made datasets including the targeted organisms (included datasets) as well as datasets including all orders, we saw in our initial study but excluding the targeted organisms (excluded datasets). The orders we saw in our initial study were: Aeromonadales, Enterobacterales, Pasteurellales, Alteromonadales, Vibrionales, Pseudomonadales, Cellvibrionales, Xanthomonadales, Steroidobacterales, Bacillales, Pseudonocardiales, Corynebacteriales, Streptosporangiales, Verrucomicrobiales, Blastocatellales, SJA-28, Nitrospirales, Rubrobacterales, and Burkholderiales.

<u>Pseudomonadales-specific designs.</u> Three unique data sets were used to design and evaluate the specificity of designs. Initially, cat-RNA were generated using a design set of 3000 Pseudomonadales, which represents the order targeted. Then, two evaluation sets were used, including an on-target evaluation set of 3000 Pseudomonadales that differed from the initial design set and an off-target evaluation set of 3000 Enterobacterales. The sequences within the design and evaluation sets for Pseudomonadales are of unique species, such that one species is not found more than once in each dataset. However, the design and evaluation sets for Pseudomonadales can have overlapping sequences between each other.

### Strains

The genotypes of the strains used are provided in Table S2. NEB Turbo competent *E. coli* (New England Biolabs) was used for cloning, *E. coli* MG1655 and *P. putida* F1 were used for evaluating cat-RNA barcoding in pure culture, *E. coli* MG1655 and *P. putida* KT2440 were used as recipient strains for conjugation, and *E. coli* MFDpir was used as a donor for conjugation.^42^ To evaluate cat-RNA barcoding in individual strains, *E. coli* MG1655 were transformed using heat shock, while *P. putida* were transformed using electroporation.^55^ For all cloning, cells were grown in LB30 medium, which contained 5 mM magnesium and was prepared by mixing 10 g tryptone, 30 g yeast extract, 10 g NaCl, and 1 g MgCl_2_·6 H_2_O in 1 L H_2_O. All other experiments used LB medium. Unless noted, *E. coli* was grown at 37°C while *P. putida* was grown at 30°C.

### Plasmids

Plasmids are described in Table S3. A plasmid (pRAM24) that transcribes a broad-host range cat-RNA was used as a universal barcoding system,^29^ which is kanamycin resistant and has a pBBR1 origin. This cat-RNA is designed to barcode U1376 in *E. coli* 16S rRNA and related locations in other rRNA. To create a plasmid that selectively targets Pseudomonadales (pPribo-B0), the guide and IGS in pRAM24 were modified using Golden Gate cloning.^56^ The barcode of pRAM24 was modified to yield a plasmid (pU1376-B2) that transcribes the universal barcoding system but uses a barcode that differs from the Pcat-RNA in pRibo-B0. Finally, a plasmid (pRAM25) that represses cat-RNA transcription by expressing CymR was used in the donor strain for conjugation experiments to repress barcoding;^29^ this plasmid confers chloramphenicol resistance. All plasmids were sequence verified and deposited in Addgene.

### RNA Extraction

Cultures of *E. coli* MG1655 or *P. putida* F1 were transformed with plasmids expressing a universal cat-RNA (pU1376-B2) and Pcat-RNA (pPribo-B0). Transformations were spread on LB-agar plates containing chloramphenicol (34 μg/mL) and kanamycin (50 μg/mL). After growing overnight, individual colonies were used to inoculate LB cultures containing chloramphenicol (34 μg/mL) and kanamycin (50 μg/mL), which were grown for 16 hours at 30°C with shaking (250 rpm). RNA was extracted using the Maxwell® Purefood GMO and Authentication Kit (Promega). Pelleted cells were resuspended in CTAB buffer (1 mL) and heated at 90°C for 5 minutes to lyse cells. Proteinase K (40 μL, 20 mg/mL) was added, mixed by vortexing, and incubated at 40°C for 10 minutes. Cell lysate (300 µL) was processed using the Maxwell® RSC 48 instrument using the Maxwell RSC PureFood GMO and Authentication Kit (Promega AS1600) and eluted using 50 μL of provided elution buffer, following the manufacturer’s protocol. The eluted samples were treated with 1 μL of TURBO DNase from the DNA-free™ DNA Removal Kit (Invitrogen, AM1906) and cleaned up using the inactivation reagent beads provided with the kit according to the manufacturer’s routine DNase treatment protocol.

### Quantitative PCR

Total transcribed cat-RNA and barcoded rRNA were measured via reverse transcriptase (RT) quantitative polymerase chain reaction (qPCR). Purified RNA was first diluted 125-fold to eliminate residual DNase. The abundance of transcribed cat-RNA was measured using primers and probes that bind within the barcode region of the cat-RNA. Primers and probes were synthesized by IDT (Table S4). All probes used a FAM fluorophore and ZEN quencher. The spliced RNA was measured using primers and probes that span the splice site of the barcoded 16S rRNA molecule. Reactions were prepared in 10 μL volumes using 4 μL of diluted RNA (diluted 125x) and the Clara Probe 1-Step Lo-ROX Mix (PCR Biosystems, PB25.81-01) was used for the reactions. Assays to measure native 16S rRNA were performed in 10 μL reactions containing 4 µL of diluted RNA (125X) and the qPCRBIO Sygreen 1-Step Go Lo-ROX mix (PCR Biosystems, PB25.31-01). Forward and reverse primers had a final concentration of 400 nM, while the probe was 200 nM. All reactions were prepared in MicroAmp™ Fast Optical 96-Well Reaction Plates, 0.1 mL (Applied Biosystems, 4346907). Reactions were carried out using a QuantStudio 5 thermocycler (Applied Biosystems). Reverse transcription was performed at 52°C for 10 minutes, followed by polymerase activation at 95°C for 3 minutes. Denaturation at 95°C for 15 seconds and annealing/extension at 62°C for 30 seconds was repeated for 50 cycles. A melt analysis was included for native 16S rRNA assays. Cq values were determined using Design and Analysis Software (Applied Biosystems, v 2.8.0).

### Conjugation

As donor strains, *E. coli* MFDpir harboring pRAM25, which encodes the CymR repressor and represses cat-RNA expression in the donor, was transformed with pU1376-B2 or pPribo-B0. Individual colonies of these donor strains were grown overnight in LB medium (3 mL) with kanamycin, chloramphenicol, and 2,6-diaminopimelic acid (DAP, 0.3 mM). These cultures were used to inoculate cultures (3 mL) containing LB medium, DAP, and antibiotics. Each culture was washed with PBS three times to remove antibiotics and resuspended at an OD of 1. *E. coli* MG1655 and *P. putida* F1 cultures prepared in a similar manner using LB medium lacking antibiotics and DAP were mixed at a 1:1 (v/v) ratio to constitute the two-member recipient community mixture. Donor cells and the recipient community mixture were then mixed at a 1:1 (v/v) ratio. Each donor-recipient mixture (50 µL) was spotted onto a nitrocellulose filter that had been placed on LB–agar medium containing DAP (0.3 mM). After incubating at 30°C for 24 hours, filters were placed in a sterile 1.5 mL tube and resuspended in 1 mL of PBS by vortexing for 10 seconds. Cells were pelleted by centrifugation (3 minutes, 6800 x g), and filters were removed. Cell pellets were stored at −80°C until RNA extraction.

### Amplicon Sequencing

Four combinations of primers were used for next-generation sequencing. For amplifying universal cat-RNA barcoded rRNA, a forward primer (oRM20) and a custom reverse primer (oRM21) that anneals to the barcode were used. For amplifying Pcat-RNA barcoded rRNA, a different forward primer (oMJD147) was selected to ensure the formation of ∼500 base pair amplicon products due to differential splicing sites between the two cat-RNAs. To amplify native 16S rRNA, reverse primers were designed for both forward primers (oRM20 and oRM18, oMJD147 and oMJD150) for amplifying a ∼500 base pair region between the V6 and V8 regions of the 16S rRNA. LunaScript® Multiplex One-Step RT-PCR Kit (NEB) was used for one-step reverse-transcription PCR reactions. Reactions (25 µL) were prepared following the manufacturer’s protocol with the supplementation of 10% DMSO. The following PCR setting were used: (i) 60 °C for 1 minute, 55 °C for 10 minutes, 98°C of 1 minute; (ii) followed by either 25 cycles (native 16S rRNA) or 40 cycles (barcoded rRNA) of 98°C for 10 seconds, 60°C for 20 seconds, and 72°C for 12 seconds; and (iii) 72°C for 5 minutes and hold at 4°C. PCR products were separated on a 2% agarose gel. Products with the expected size were then gel-purified, purified using the Monarch DNA Gel Extraction Kit Protocol (NEB #T1020), and sequenced using Amplicon-EZ (Azenta).

### NGS Analysis Pipeline

Data analysis of sequenced amplicons was performed as described with modifications in trimming.^29^ All raw reads were imported to Qiime2 (version 2024.2). The Cutadapt plugin was used for trimming raw reads of primer sequences and cat-RNA barcodes, using the default allowed error rate of 10%. For Universal cat-RNA treated samples’ native rRNA amplicons, forward primer (GCAACGCGAAGAACCTTACC) and reverse primer (TGACGGGCGGTG<u>W</u>GT<u>R</u>CA) were removed. For universal cat-RNA barcoded rRNA amplicons, the forward primer (GCAACGCGAAGAACCTTACC), reverse primer (AAGTCATGCCGTTTCATGTGATC), and barcode (CGGATAACCACTCGGGAAAAGCATTGAACACCAT) were removed. For Pcat-RNA treated samples’ native rRNA amplicons, the forward primer (GGATTAGATACCCTGGTAGTC) and reverse primer (CATTGTAGCACGTGTGTAGC) were removed. For Pcat-RNA barcoded rRNA amplicons, the forward primer (GGATTAGATACCCTGGTAGTC), reverse primer (AAGTCATGCCGTTTCATGTGATC), and barcode (CGGATAACGGGAAAAGCATTGAACACCAT) were removed. The DADA2 plugin was used to denoise and dereplicate reads. The resulting feature tables were filtered to retain ASVs with a representative sequence of at least 250 base pairs. Additionally, only ASVs present in at least two out of the three biological replicates and with at least three reads were kept for downstream analysis. Taxonomy was assigned using feature-classifier classify-sklearn using the pre-formatted SILVA 138 SSURef NR99 full-length classifier. All ASVs classified under Enterobacterales were considered as *E. coli*, and all ASVs classified under Pseudomonadales were considered as *P. putida*. All other ASVs (on average 1.23% of reads per sample) were discarded.

### Statistical Analysis

Taxonomy-assigned read counts were rarefied to 100,000 reads prior to any statistical analysis. Pair and unpaired two-tailed Welch’s T-tests were used to evaluate differences between rarefied ASV read counts. All statistical analyses were performed in RStudio (version 2023.12.1).

## Author contributions

K.R.G, L.B.S, and J.J.S: Conceptualization; K.R.G, T.S., L.F., and M.J.D: Investigation; K.R.G, T.S., L.F., and M.J.D: Analysis; K.R.G, T.S., L.F., M.J.D, L.B.S, and J.J.S: Drafting manuscript; K.R.G, T.S., L.F., M.J.D, J.C., L.B.S, and J.J.S: Manuscript review and editing; K.R.G, T.S., L.F., M.J.D, L.B.S, and J.J.S: Visualization; J.C., L.B.S, and J.J.S: Funding; L.B.S and J.J.S: Supervision.

## SUPPORTING INFORMATION

Supplementary figures and tables referenced throughout the main text, providing additional analysis of archaeal and eukaryote designs, more details on next-generation sequencing from conjugation measurements, and primers and probes used for analysis.

