## Supplemental Figures for "Controlling the taxonomic composition of biological information storage in 16S ribosomal RNA"

### TABLE OF CONTENTS


| Item | Title | Page |
|---|---|---|
| Fig. S1 | Uracil conservation across diverse archaeal 16S rRNA | 3 |
| Fig. S2 | Uracil conservation across diverse eukaryotic 18S rRNA | 4 |
| Fig. S3 | Guide sequence optimization by ambiguity reduction | 5 |
| Fig. S4 | Comparing cat-RNA designs for archaea | 6 |
| Fig. S5 | Comparing cat-RNA designs for eukaryotes | 7 |
| Fig. S6 | Analysis of 16S rRNA levels in bacterial samples | 8 |
| Fig. S7 | 16S rRNA NGS data before rarefaction | 9 |
| Fig. S8 | Barcoded-rRNA NGS data before rarefaction | 10 |

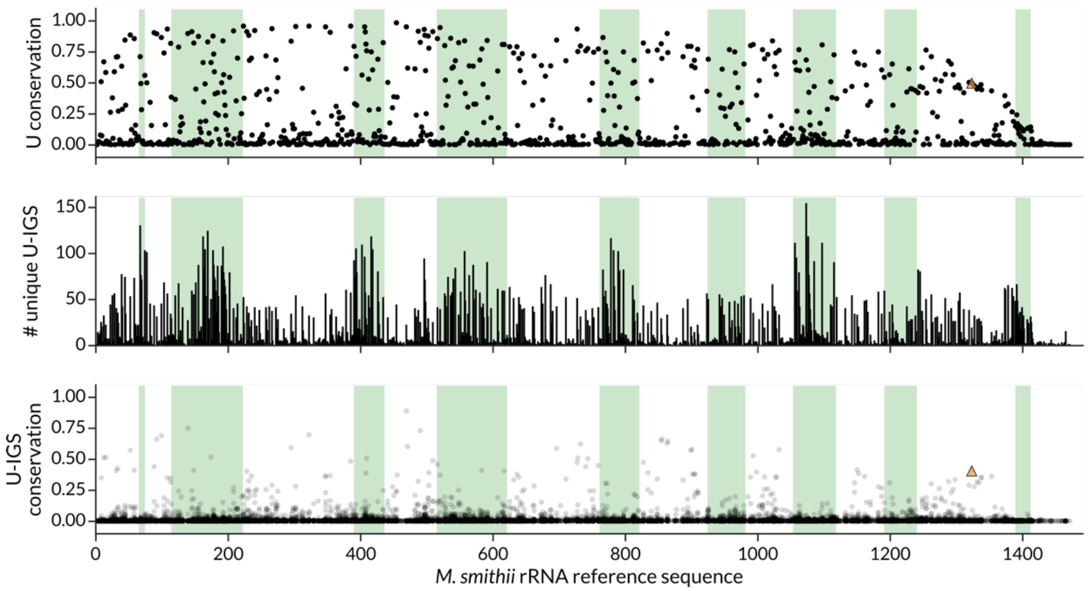


**Figure S1. Uracil conservation across diverse archaeal 16S rRNA.** (*top*) The frequency of U at each position in the reference 16S rRNA, (*middle*) the diversity of IGS-binding pentanucleotide sequences adjacent to each U, and *(bottom)* the frequency of each U-pentanucleotide sequence. The archaeal 16S rRNA test set included 337 sequences, and *Methanobrevibacter smithii* was used as a reference sequence. The 16S rRNA hypervariable regions, indexed to the reference sequence (*M. smithii*), are in green.

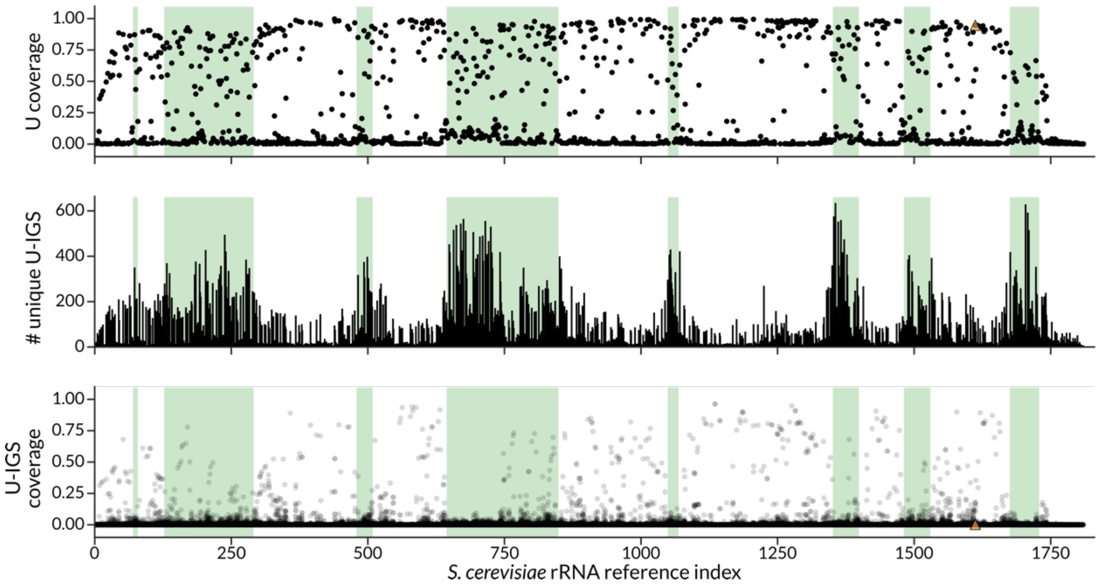


**Figure S2. Uracil conservation across diverse eukaryotic 18S rRNA.** (*top*) The frequency of U at each position in the reference 18S rRNA, (*middle*) the diversity of IGS-binding pentanucleotide sequences adjacent to each U, and (*bottom*) the frequency of each U-pentanucleotide sequence. The eukaryotic 18S rRNA test set included 2618 sequences, and *Saccharomyces cerevisiae* was used as a reference sequence. The 18S rRNA hypervariable regions, indexed to the reference sequence (*S. cerevisiae*), are in green.

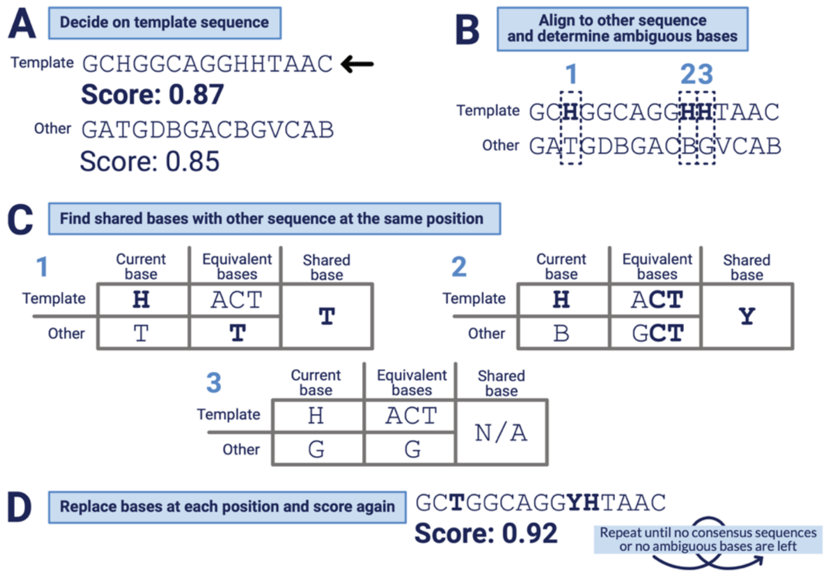


**Figure S3. Guide sequence optimization by ambiguity reduction.** Ribodesigner optimizes the guide sequence design by using the other consensus sequences generated. (**a**) It first finds the highest scoring sequence and names it the template sequence, and picks the next highest scoring as the ”other” sequence. (**b**) Then, it finds any position in the template with ambiguity bases to optimize and finds the corresponding position on the other sequence. (**c**) At each position, Ribodesigner determines if there are any shared bases that would reduce the ambiguity. (**d**) If any are found, Ribodesigner replaces the ambiguous base with the shared base and scores the resulting guide again. This process is repeated until either there are no more consensus sequences to compare against, or until there are no ambiguous bases left in the template.

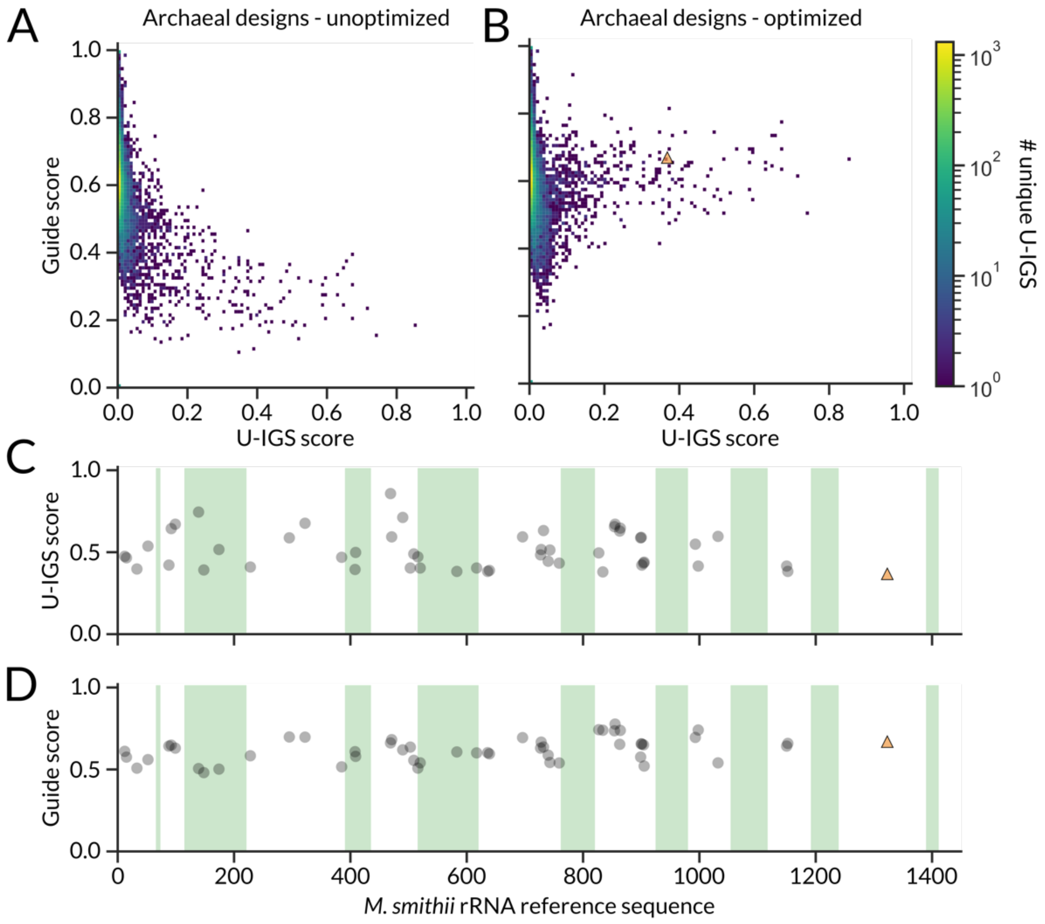


**Figure S4. Comparing cat-RNA designs for archaea.** (**A**) Comparison of guide designs that were generated by evaluating small batches (n = 10). (**B**) Comparison of the guide designs that were generated by combining the winners from each batch to further optimize guide designs. (**C**) For the 50 designs having the highest U-IGS scores, the location of U targeted for barcoding is mapped onto *M. smithii* rRNA. (**D**) For those same designs, the guide score*s* are also shown. The cat-RNA previously applied in a wastewater is shown as an orange triangle.

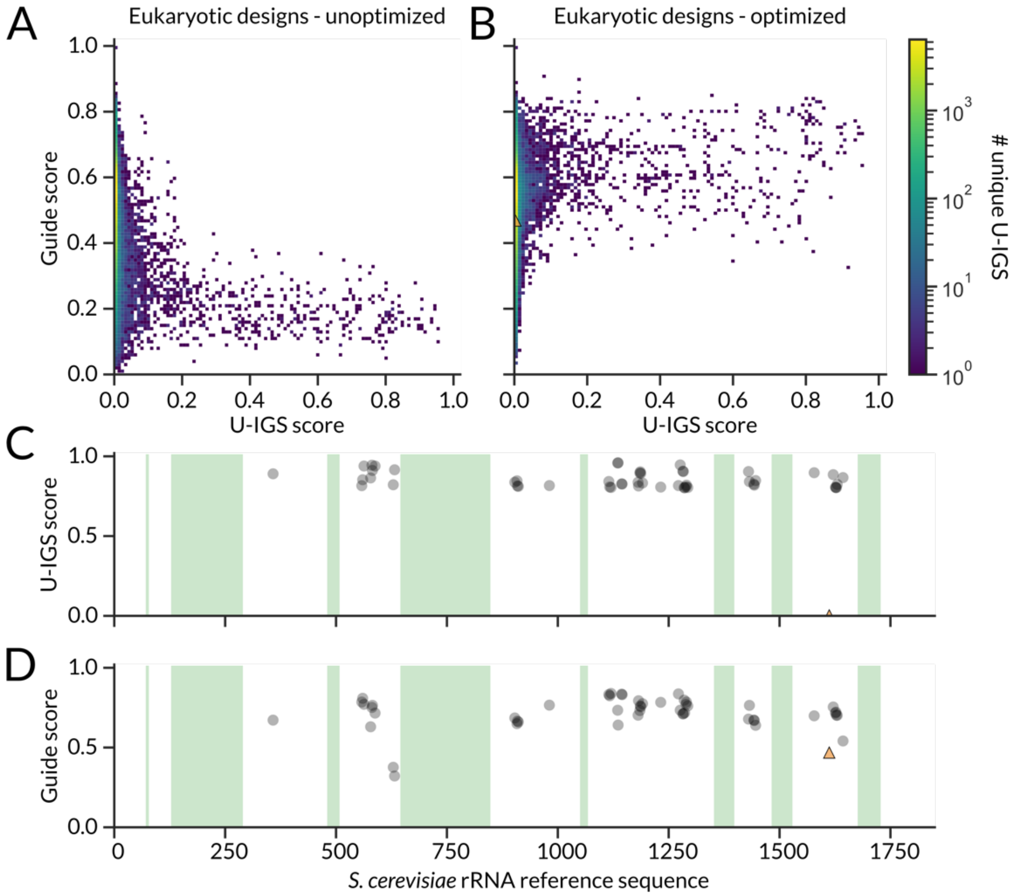


**Figure S5. Comparing cat-RNA designs for eukaryotes.** (**A**) Comparison of guide designs that were generated by evaluating small batches (n = 10). (**B**) Comparison of the guide designs that were generated by combining the winners from each batch to further optimize guide designs. (**C**) For the 50 designs having the highest U-IGS scores, the location of U targeted for barcoding is mapped onto *S. cerevisiae* rRNA. (**D**) For those same designs, the guide scores are also shown. The cat-RNA previously applied in a wastewater is shown as an orange triangle.

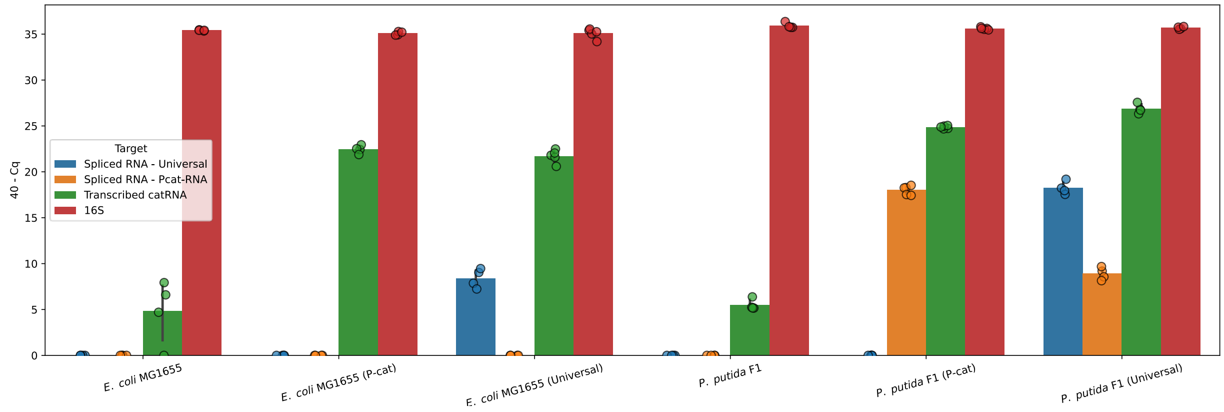


**Figure S6. Analysis of 16S rRNA levels in bacterial samples.** To evaluate the relative amount of 16S rRNA in samples analyzed for splicing, qPCR was used to evaluate total 16S rRNA (red) in *E. coli* MG1655 and *P. putida* F1 lacking or containing different vectors that transcribe the universal cat-RNA and Pcat-RNA. Additionally, qPCR was used to quantify the amount of cat-RNA and Pcat-RNA transcribed (green), the amount of 16S rRNA that was barcoded by the universal cat-RNA (blue), and the amount of 16S rRNA that was barcoded by Pcat-RNA (orange).

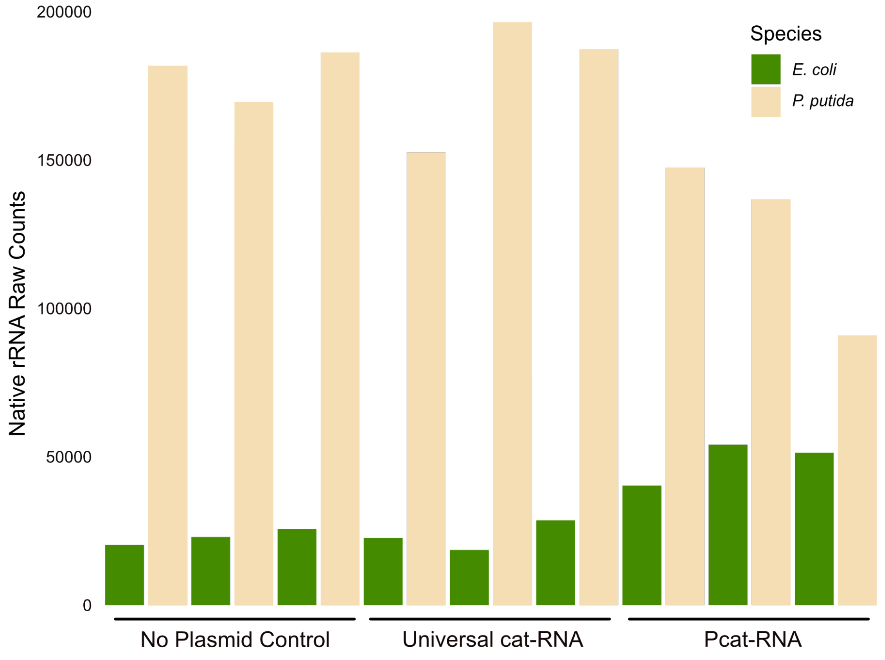


**Figure S7. 16S rRNA NGS data before rarefaction**. Each bar represents the raw count of 16S rRNA reads from different biological replicates from conjugation reactions that contained *E. coli* MFDpir*, E. coli* MG1655, *P. putida* KT2440 at ratios of 2:1:1. When analyzing the control and broad-host range cat-RNA (universal), primers were used that generated an amplicon that included variable regions V6-V8. In contrast, when analyzing Pcat-RNA samples, primers were used that generated an amplicon that included variable regions V5-V6.

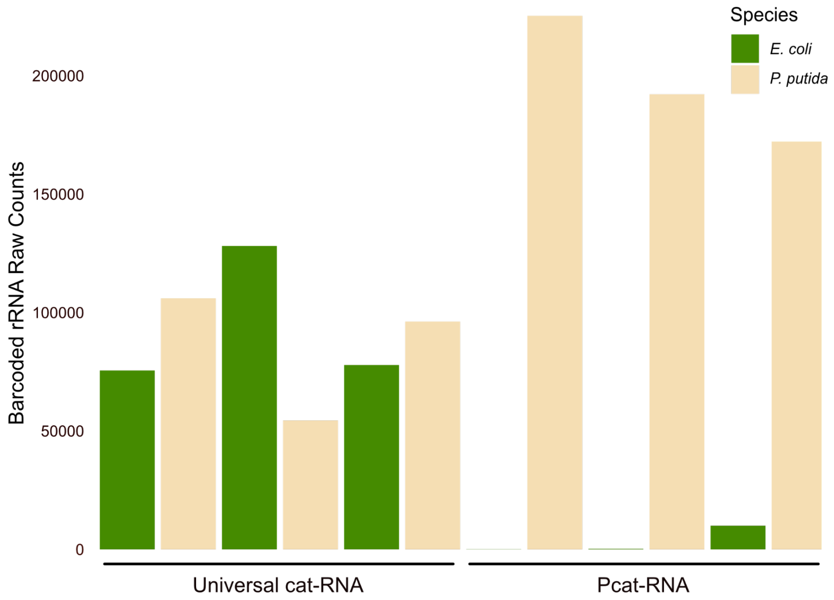


**Figure S8. Barcoded-rRNA NGS data before rarefaction**. Each bar represents the raw count of barcoded-rRNA reads from different biological replicates from conjugation reactions that contained *E. coli* MFDpir*, E. coli* MG1655, *P. putida* KT2440. The variable regions V6-V8 of the 16S rRNA were targeted for sequencing with the universal cat-RNA and the V5-V6 regions were targeted with the Pcat-RNA samples as the P-cat-RNA adds a barcode upstream of the V7 region.
